# The road to restoration: Identifying and conserving the adaptive legacy of American chestnut

**DOI:** 10.1101/2023.05.30.542850

**Authors:** Alexander M Sandercock, Jared W Westbrook, Qian Zhang, Jason A Holliday

## Abstract

The American chestnut (*Castanea dentata*) is a functionally extinct tree species that was decimated by an invasive fungal pathogen in the early 20^th^ century. Disease resistant chestnuts have been developed through hybridization and genetic modification, but these populations may lack the adaptive genomic diversity necessary to restore the species across its climatically diverse historical range. An understanding of the genomic architecture of local adaptation in wild American chestnut and identification of seed zones for germplasm conservation are necessary in order to deploy locally adapted, disease-resistant American chestnut populations. Here, we characterize the genomic basis of climate adaptation in remnant wild American chestnut, define seed zones based on climate envelopes and adaptive diversity, and make sampling recommendations for germplasm conservation. Whole genome re-sequencing of 384 trees coupled with genotype-environment association methods suggest the species range can be most parsimoniously subdivided into three seed zones characterized by relatively homogeneous allele frequencies relative to rangewide adaptive diversity. Using these data, we developed a method to estimate the number of samples required from each seed zone to recapitulate standing adaptive diversity in each seed zone, and found that on average, 21-29 trees will need to be conserved to capture 95% of the wild adaptive diversity. Taken together, these results will inform the development of an *ex situ* germplasm conservation and breeding plan to develop locally adapted blight-resistant American chestnut populations, and provide a blueprint for developing restoration plans for other imperiled tree species.

## 1 Introduction

American chestnut (*Castanea dentata*) is one of the tree species most vulnerable to climate change in the United States (Potter et al., 2017). A large deciduous tree native to the Appalachian Mountains of eastern North America, the American chestnut was decimated by a fungal blight beginning in the early 20th century that killed billions of trees. While approximately 400 million chestnut stems remain throughout the range, these are almost exclusively root collar sprouts that rarely achieve more than a few years of growth before being reinfected with blight (Paillet, 2002; Scrivani, 2011). This repetitive cycle of growth and die-back and the lack of widespread reproduction makes American chestnut functionally extinct in the wild (Paillet, 2002).

Efforts to develop blight resistant American chestnut through introgression from Chinese chestnut (*Castanea mollissima*) (Steiner et al., 2017) and insertion of an oxalate oxidase (OxO) transgene may soon enable widespread restoration plantings (Newhouse et al., 2014; Powell et al., 2019; Zhang et al., 2013). However, particularly in the case of the genetically-modified lines, a lack of genetic diversity in general, and adaptive diversity in particular, necessitates an outcrossing strategy to both increase effective population size and target specific families to the climate of local planting sites. And while the hybridization and backcrossing program has incorporated some standing variation via pollination from wild trees, the extent of adaptive variation segregating in backcross populations is unknown.

Adaptation to local climate in temperate forest trees is determined by multiple abiotic constraints, including the correct timing of growth and dormancy transitions and associated cold acclimation, maximum winter cold hardiness, water-use efficiency, and drought tolerance or avoidance (Howe et al., 2003; Savolainen et al., 2007). Similar to many temperate tree species, American chestnut has migrated from glacial refugia to occupy extant ranges spanning thousands of kilometers from southern to northern limits (Sandercock et al., 2022). Differentiation in locally-adaptive traits related to temperature variation has been documented in common gardens(Saielli et al., 2012, 2014; Schaberg et al., 2022), which suggest adaptive clines similar to other widespread temperate tree species. However, because of the rarity of wild reproduction, it is exceptionally difficult or impossible to assemble chestnut common gardens that encompass locally dense and geographically widespread sampling of the historical range. Hence, identifying the genomic targets of climate-related selection is a key step in understanding the underlying mechanism of local adaptation in this species, as well as for developing strategies for capturing and incorporating the adaptive legacy of the species. One way to approach this objective is through the concept of seed zones, whereby a species range is divided into the smallest number of regions that effectively describe how climate and ecoregions are partitioned. Ideally these zones also reflect reaction norms from reciprocal common gardens, although such data is usually only available for species of high economic value. Existing climate and ecological data suggest >10 different seed zones within the historical American chestnut range (Bower et al., 2014; Pike et al., 2020), but these are not specific to chestnut, and may or may not reflect how adaptive traits and genomic diversity is arrayed across the landscape for this species. Similarly, these zones do not inform the sampling intensity required per zone to capture standing adaptive diversity.

Here, we describe the genomic basis of local adaptation in American chestnuts and make sampling recommendations to inform germplasm conservation efforts. We leverage a *≈*21 million SNP whole-genome sequencing (WGS) dataset developed by Sandercock et al. (2022) to (i) identify loci significantly associated with climate; (ii) define seed zones for germplasm conservation; and (iii) develop a novel method for making sampling recommendations to capture wild adaptive diversity. These results will inform American chestnut restoration efforts, and provide a blueprint for similar conservation efforts in the future.

## 2 Methods

### 2.1 Dataset preparation

We used the 21,136,994 filtered SNPs and 356 *C. dentata* individuals that were previously reported by Sandercock et al. (2022) (Figure 1). Briefly, 384 leaf samples were obtained from throughout the American chestnut natural range and then their DNA was sequenced using whole-genome sequencing (WGS) at HudsonAlpha Institute for Biotechnology. After filtering the 384 individuals based on missing data and ancestry tests, 356 *C. dentata* samples were included in the final dataset. Missing sites were imputed using BEAGLE v5.4 (Browning et al., 2018) and a minor-allele frequency (MAF) filter was applied using PLINK v1.9 to remove SNPs with MAF < 0.05 (Purcell et al., 2007). The final genomic dataset contained 11,526,713 SNPs and 356 samples.

**Figure 1:**
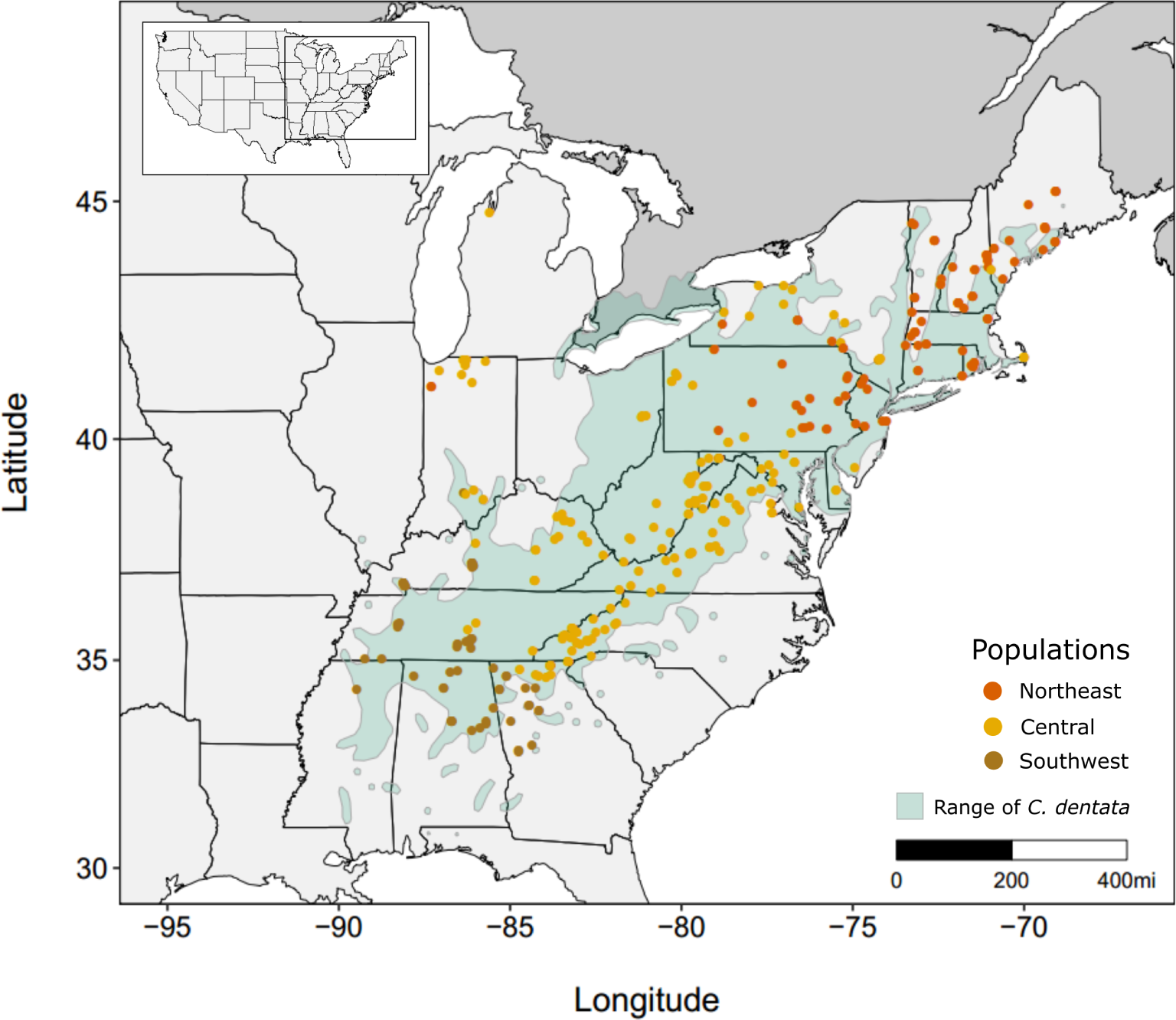
Range map of *Castanea dentata* and the locations of the 356 American chestnut samples and their assigned population from Sandercock et al. (2022).

Environmental data was obtained from climateNA (Wang et al., 2016) and comprised 32 monthly and seasonal climate variables (Table S1). The environmental variables were the average for 1961-1990 and the ensemble projections under the CMIP6 future climate scenarios for 2050 and 2080 (AdaptWest Project, 2022; Mahony et al., 2022). Climate data was not available for three of the samples, so they were removed from the analysis. Collinearity between climate variables was assessed to exclude highly correlated variables that could increase false-positive rates in genotype-environment relationships (Capblancq & Forester, 2021). We first used gradientForest to rank the importance of each of the climate variables using a random subset of 50k SNPs (Ellis et al., 2012; Fitzpatrick & Keller, 2015). Pearson’s correlations between each pair of climate variables were estimated using the pairs.panel() function from the *psych* R package (Revelle, 2022). Climate variables were removed if they were highly correlated (R^2^>0.7) with another variable of higher importance from the gradientForest predictor importance results.

### 2.2 Identifying climate-associated adaptive loci in American chestnut populations

Univariate and multivariate genotype-environment association (GEA) tests were performed to identify putatively adaptive loci that were associated with climate. Redundancy analysis (RDA) is a multivariate genotype-environment association method (GEA) that accounts for all environmental variation simultaneously. RDA has been shown to have higher true positive and lower false positive rates than univariate and several other multivariate approaches (Forester et al., 2018), and enables assessment of the contributions of multiple factors (climate, geography, and population structure) to patterns of genetic variation. For the RDA analysis, we used the vegan R package (Oksanen et al., 2020). First, we used a partial RDA (pRDA) to evaluate the contributions of each of climate, geography, and population structure to rangewide diversity. Specifically, we performed four tests: a full model with all factors, a climate only model, a geography only model, and a population structure only model (Capblancq & Forester, 2021), and summarized the influence of each factor on genetic variation with an ANOVA in R. For the pRDA analyses, we used a linkage-disequilibrium (LD) pruned dataset due to computational efficiency. An LD-prune was applied to the *≈*11.5 million SNP dataset using PLINK v1.9 to include SNPs with R^2^ values <0.5 within 50 SNP sliding windows (step size 10 SNPs) – resulting in a reduced dataset of 2,934,239 SNPs (Purcell et al., 2007). We then performed a GEA scan of the *≈*11.5M SNP dataset and the 10 uncorrelated climate variables following Capblancq and Forester (2021). As correcting for population structure can cause the RDA outlier tests to be overly conservative, we report only the uncorrected model (Forester et al., 2018; Lotterhos, 2022). We retained three RDA axes, and the top 0.1% of p-values were identified as candidate locally adaptive sites.

As a complementary method to detect genotype-environment relationships, we used latent factor mixed models (LFMM) as implemented in the R package LEA (Caye et al., 2019; Frichot & Francois, 2015). LFMM2 uses univariate regression to identify linear associations between loci and climatic variables, while accounting for confounding factors such as background allelic covariance arising from population structure(Caye et al., 2019). We first summarized the 10 climatic variables with PCA, retaining the first three PC axes, which explained *≈*89% of variance within the climate variables dataset (Oksanen et al., 2020). The lfmm2() function in the LEA package in R was used to run LFMM2 three times (once for each environmental PC axis) with K=2 latent factors to correct for population structure (Frichot & Francois, 2015). P-values, z-scores, and the genomic-inflation factors (GIF) were estimated using the lfmm2.test() function with default parameters (Frichot & Francois, 2015). To account for multiple testing, p-values were converted to q-values using the qvalue() function in R (Storey et al., 2021). Loci that had a q-value < 0.1 (FDR < 10%) were retained as candidate adaptive loci. Finally, outlier loci identified by the RDA and the three LFMM runs were combined for subsequent analyses.

To assess whether the putatively adaptive loci were enriched for coding or noncoding regions of the genome, the locations of the outlier SNPs were cross-referenced with the American chestnut genome feature file (Cdentata_673_v1.1.gene_exons.gff3.gz; http://phytozome-next.jgi.doe.gov/). Introns were not included in the GFF file, and were added using the ATAG biopython package (Dainat, 2022). Promoters were also added to the GFF file as the 2 kb region upstream of the mRNA intervals.

### 2.3 Spatial distribution of adaptive genomic diversity and genomic offset

To characterize the spatial distribution of adaptive genomic variation throughout the chestnut range, we used the Gradient Forest (GF) machine learning algorithm(Fitzpatrick & Keller, 2015). GF estimates nonlinear turnover functions of allele frequencies across environmental gradients. Uniquely, the shape of the nonlinear turnover functions estimated by GF enable rescaling of the environmental gradients to reflect the specific ways in which the multidimensional genomic variation is constrained by the climate space. To do this, we first estimated a gradientForest model using the 18,483 adaptive SNPs from the LFMM and RDA analyses as response, and the climate information from the 10 uncorrelated climate variables for each sampling location as the predictors. This trained model was then used to predict the spatial distribution of genomic variation over the entire chestnut range. A PCA was then performed on the GF model prediction and the top three principal components (PCs) were used to visualize the multidimensional adaptive genomic variation across the American chestnut range. To understand how climate change may impact the relationship between genome and environment, we performed the same procedure outlined above using two future climate predictions for 2080 (RCP4.5 and RCP8.5). We then calculated the Euclidean distance between the current climate GF model and the future climate model, which reflects ‘genomic offset’ – an estimate of future maladaptiveness due to climate change.

### 2.4 Identifying seed zones within the American chestnut natural range

A major objective of this research was to partition the chestnut range into geographic seed zones that reflect relatively homogeneous areas with respect to multivariate adaptive genomic variation, which can subsequently be used to both conserve germplasm *ex situ*, and guide restoration plantings of blight-resistant chestnut families/lines with similar genomic compositions. To do this, the PCA loadings from the GF model above were clustered using the Optimal_Clusters_KMeans() function from the ClusterR package in R to determine the optimal number of seed zones (Mouselimis, 2022; Yu et al., 2022). Between K=1 and K=12 clusters were evaluated for fit to the data using AIC, sum of within-cluster-sum-of-squares-of-all-clusters (WCSSE), variance explained, and BIC. We then used the clara() function from the ClusterR package with K = 3 (Maechler et al., 2021); the resulting clusters are the seed zones – representing a genetically similar region relative to its adaptive variation.

### 2.5 Estimating the number of trees to sample to capture wild adaptive diversity for germplasm conservation

To estimate the number of trees required to sample from each seed zone to recapitulate the multivariate allele frequencies at adaptive loci and inform germplasm conservation, we created a custom Python script. The Python script takes the input of a VCF file and optionally a list of SNP and sample information, and performs bootstrap re-sampling of the dataset to compare allele frequencies (AF) for all SNPs from the bootstrap samples with those of the full population using linear regression. Bootstrap sampling of the full population continues until the R^2^ score from the linear regression meets or exceeds the user-defined R^2^ score (a proxy for the percent diversity to capture), then the bootstrap sampling ends and the number of subsampled trees is recorded. The process is then repeated for a predefined number of iterations (in our case, 100), and the number of trees to sample is averaged across all iterations. The final output is the mean number of trees to sample from a population to meet the desired coefficient of determination between the adaptive multivariate AFs of the sample and that of the full dataset for each seed zone.

## 3 Results

### 3.1 Drivers of genomic variation across the range of American chestnut

We previously showed that the American chestnut range can be divided into three populations with longitudinal boundaries that roughly divide the species into southern, central, and northern groups (Sandercock et al., 2022). As these boundaries also track temperature and precipitation gradients, we predicted that population structure and climate would have the largest influence on genomic variation. The partial RDA models show that climate had the largest proportion of explainable variance (0.38), followed by population structure (0.13), and geography (0.08), and that all pRDA models were significant (p<0.001,Table 1). The full pRDA model accounted for 11% of the total variance.

**Table 1:** Drivers of genomic variation in American chestnut. Four partial RDA models were performed to evaluate the influence that climate (10 environmental variables), population structure (first two PC axes from PCA on genomic dataset), and geography (geographic coordinates of each sample) had on the patterns of genomic variation throughout the American chestnut natural range. An ANOVA was performed after each partial RDA to estimate statistical significance.

| Partial RDA models | Inertia | Adj. $R^2$ | p ($>F$) | Proportion of explainable variance | Proportion of total variance |
| --- | --- | --- | --- | --- | --- |
| Full model | 79190 | 0.063 | 0.001*** | 1 | 0.11 |
| Pure climate | 30117 | 0.012 | 0.001*** | 0.38 | 0.04 |
| Pure structure | 10018 | 0.008 | 0.001*** | 0.13 | 0.01 |
| Pure geography | 6368 | 0.003 | 0.001*** | 0.08 | 0.01 |
| Confounded climate/structure/geography | 32687 |  |  | 0.41 | 0.05 |
| Total unexplained | 632195 |  |  |  | 0.89 |
| Total inertia | 711385 |  |  |  | 1.00 |

\*\*\* $p \leq 0.001$

We identified 10 uncorrelated climate variables to test for genotype-environment associations: **RH** (mean annual relative humidity (%)), **Eref** (Hargreaves reference evaporation (mm)), **PPT_wt** (winter precipitation (mm)), **EXT** (extreme maximum temp. over 30 years (°C)), **TD** (temp. difference between mean warmest month and mean coldest month (°C)), **MAR** (mean annual solar radiation (MJ m-2 d-1)), **PPT_sm** (summer precipitation (mm)), **CMI** (Hogg’s climate moisture index (mm)), **AHM** (annual heat-moisture index (mean annual temp.+10)/(mean annual precip./1000)), and **DD_18** (degree-days below 18°C) (Figure 2a, Figure S1). Since LFMM is a univariate method, the LFMM analyses utilized three PC axes as synthetic predictors in place of the 10 environmental variables due to computational demand and to reduce the number of corrections needed to account for multiple testing. For LFMM, we identified 4,988 SNPs associated with PC1(MAR, PPT_wt, TD), 2,923 SNPs associated with PC2(AHM, CMI, EXT), and 0 SNPs associated with PC3(RH) (Table S2). In total, LFMM identified 7,911 outlier SNPs (FDR<0.1) and the RDA analysis revealed 11,526 outlier SNPs (0.1% outliers), with 954 of these SNPs shared between the two GEA methods (Figure S5), for a combined total of 18,483 putatively adaptive SNPs.

**Figure 2:**
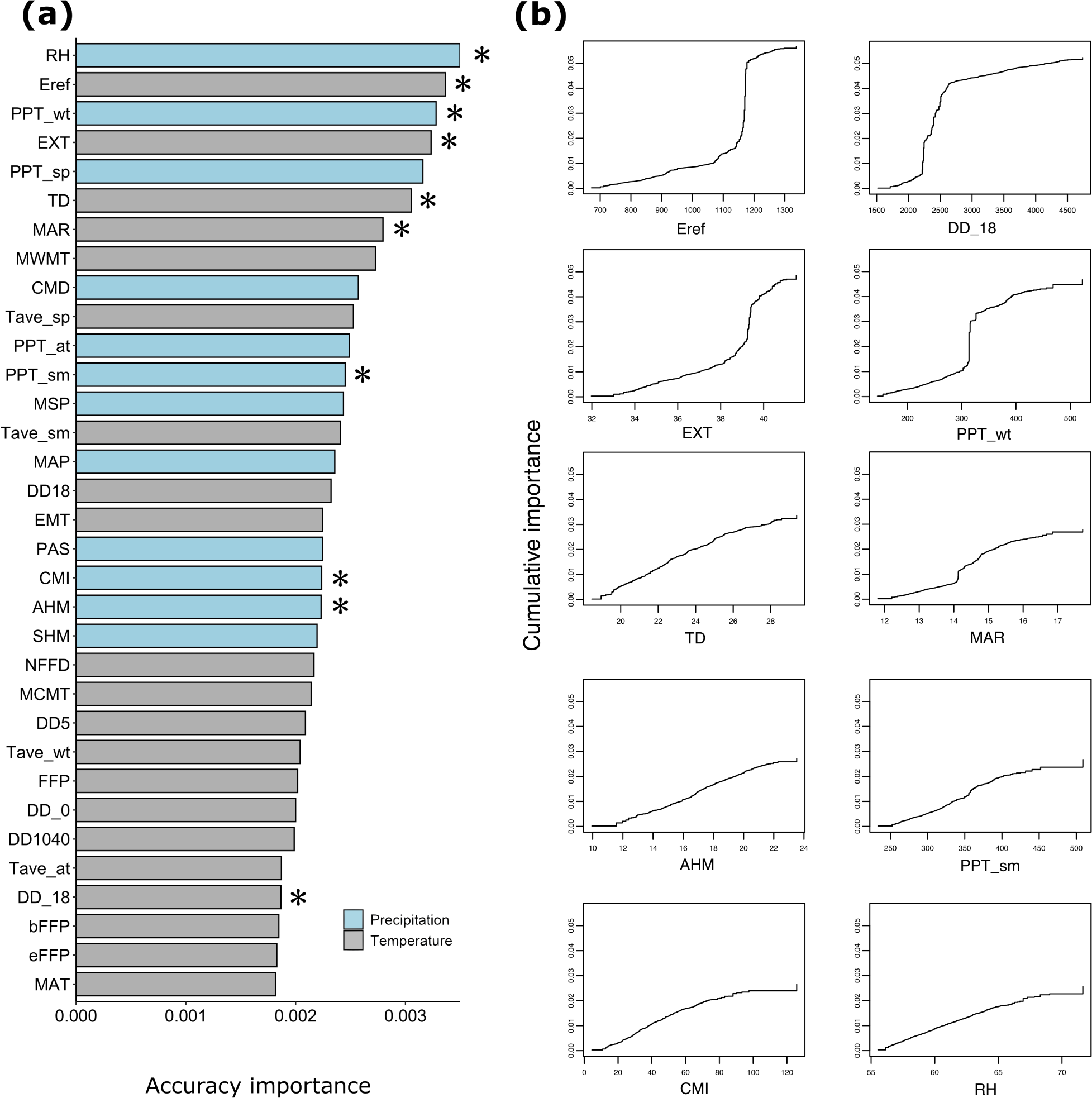
Importance of climatic variables in patterns of allelic turnover for wild American chestnut populations. A) Ranked accuracy importance of 33 climateNA variables from a gradientForest analysis that included 353 American chestnut samples and 50k SNPs. The variables denoted with ‘*’ were used for the analyses. B) Predictor cumulative importance plot. Cumulative change in allele composition across the 10 climate gradients and the 18,483 putatively adaptive loci. Sharp increases suggest a point in the climate gradient where allelic composition changes.

Among climate variables, allelic turnover was relatively linear for temperature differentials (TD), annual heat:moisture indices (AHM), summer precipitation (PPT_sm), Hogg’s climate:moisture indices (CMI), and mean annual relative humidity (RH) (Figure 2b). By contrast, sharp turnovers occurred at evaporative demand (Eref) between 1150 and 1200 mm, extreme temperatures (EXT) between 39 and 40 °C, cooling degree-days (DD_18) between 2000 and 2500 days, and winter precipitation (PPT_wt) between 300 and 350 mm (Figure 2b). The remaining climate variable, mean annual solar radiation (MAR), showed a slight increase in allele turnover rate between 14 and 15 (MJ m-2 d-1) (Figure 2b).

Throughout the genome, 210 genes contained at least one adaptive site, with most of the ‘adaptive’ genes located on chromosome one, five, and 10 (Figure S3). Approximately 18.6% of the putatively adaptive SNPs were located in promoter and gene regions, which was lower than in the random dataset (*≈*28.6%), and the initial filtered dataset (*≈*28.3%) (Table S3).

### 3.2 Geographic distribution of genomic variation and maladaptation under current and future climates

Our genotype-environment analysis suggests differences in the climatic drivers of adaptive genomic variation between northern and southern portions of the American chestnut range. The northern portion of the range was associated with temperature and precipitation (TD, DD_18, AHM), while the southern portion of the range was associated with evaporative demand and extreme temperatures (Eref and EXT) (Figure 3a,b). An additional area in western North Carolina and central portions of the Appalachian Mountains was associated with precipitation and solar radiation (CMI, MAR, PPT_sm, and PPT_wt) (Figure 3a,b). To understand how climate change will impact the match between current and future adaptive matches, we estimated genomic offset for 2080 for both middle and high emissions climate scenarios (RCP4.5 and RCP8.5, respectively). Under both climate change scenarios, the central portions of the American chestnut range were predicted to experience elevated maladaptation, effects that were exaggerated for the high emissions scenario (Figure 3c).

**Figure 3:**
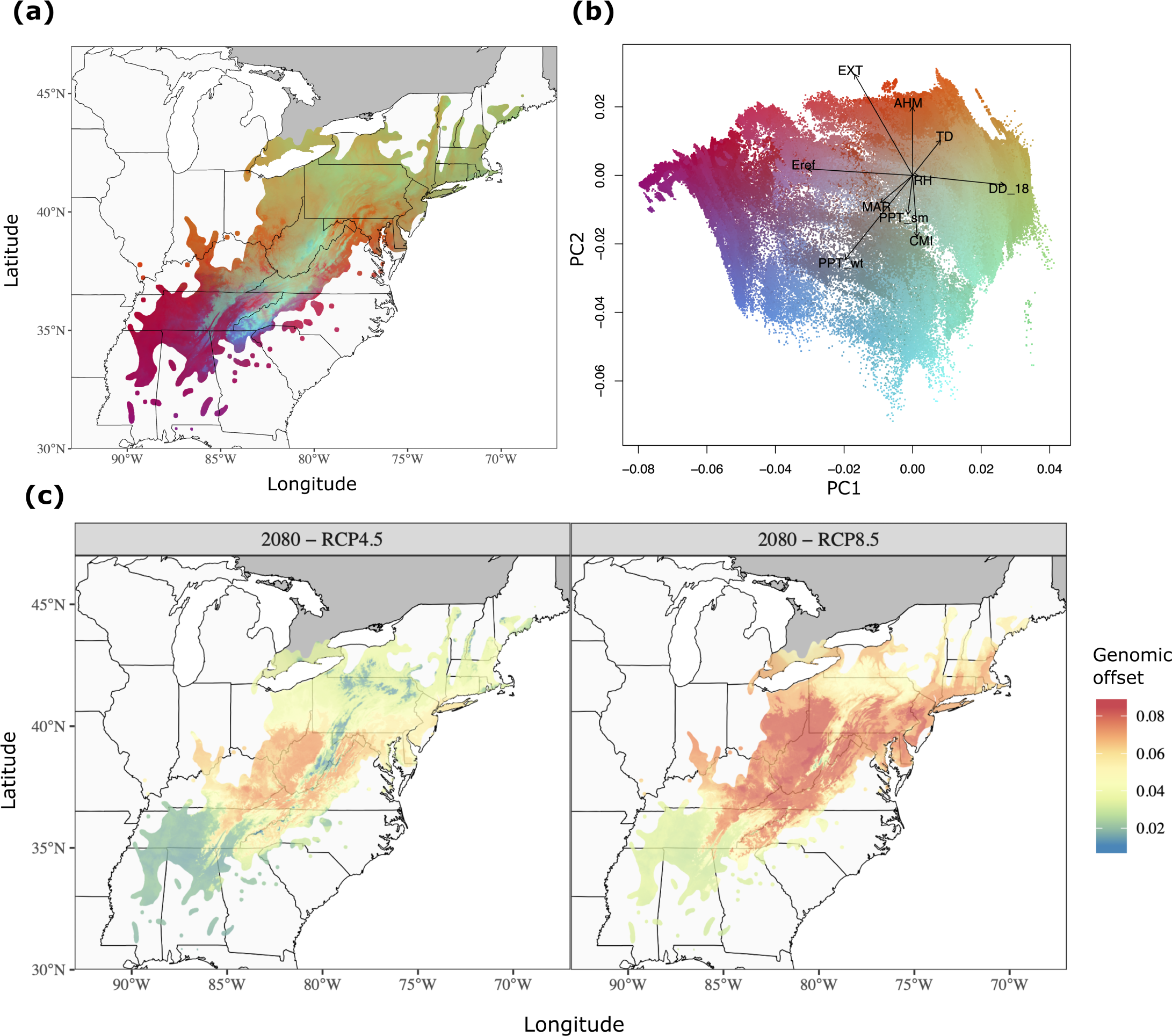
Adaptive genomic diversity under past and future climate conditions. (a) PC loadings from the gradientForest model of 10 uncorrelated climate variables and 18,483 putatively adaptive loci. (b) Biplot of loadings for PC1 and PC2. (c) Genomic offset under 2080 climate projections for moderate emissions (RCP4.5; left) and severe emissions (RCP8.5; right) scenarios.

### 3.3 Identifying seed zones

As climate was the greatest driver of genomic variation in American chestnut (Table 1), we hypothesized that we could partition the range into seed zones with boundaries reflecting precipitation and temperature gradients in the eastern US. GradientForest model selection from K=1-12 climate-based seed zones suggested that two or three were the most likely number of zones (Figure S6,Figure S7,Figure S8,Figure S9). With three such zones, the range was split into north, central, and south regions, with latitudinal boundaries at approximately 36N and 39N (Figure 4a,b). Genetic differentiation between seed zones was approximately 5.5-6.5 fold higher at adaptive loci compared with neutral loci (Figure 4c).

**Figure 4:**
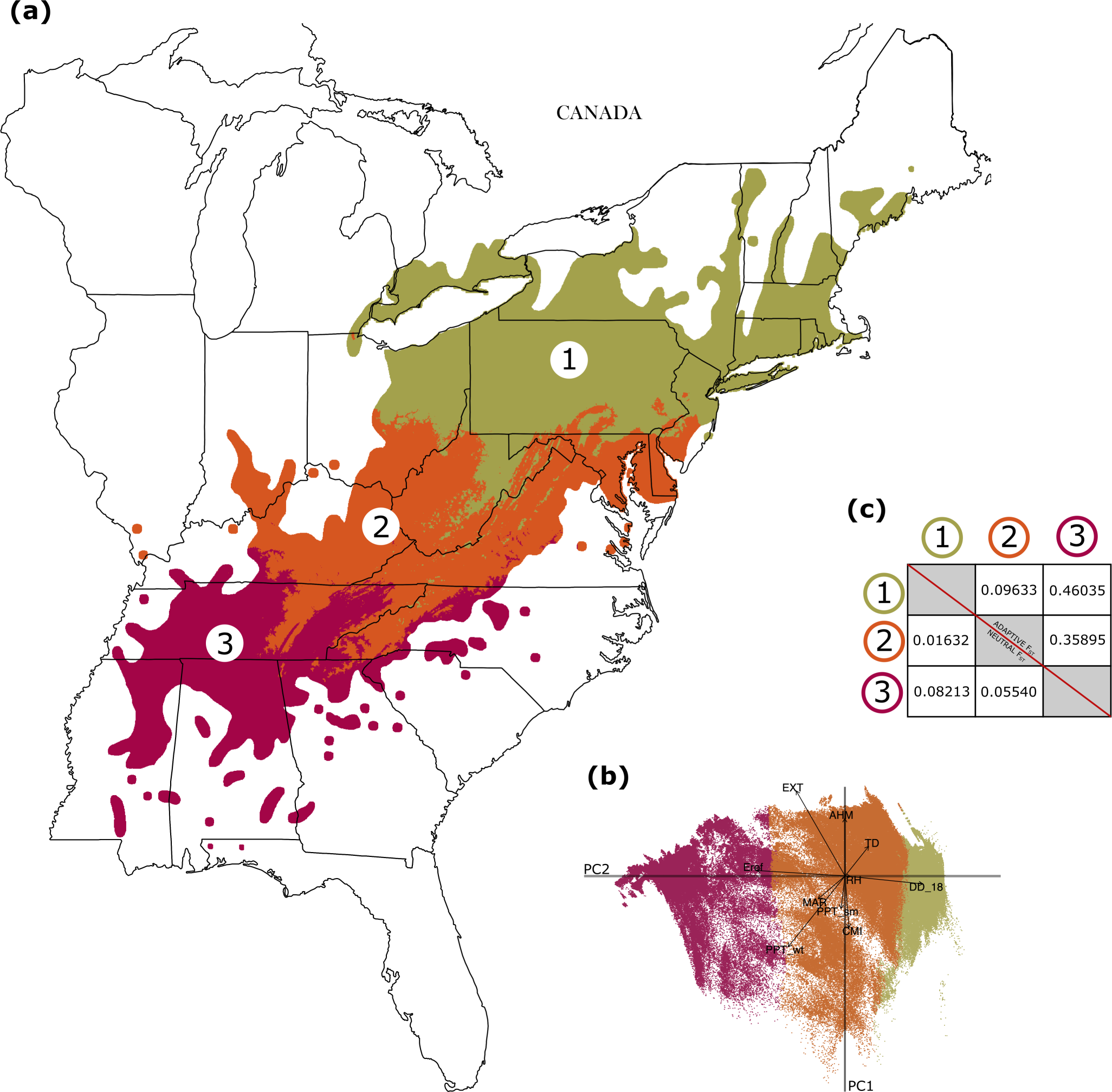
Three seed zones for American chestnut conservation that are based on the GF-predicted genomic variation. (a) Spatial distribution of the three seed zones. (b) PCA biplot of the GF-predicted genomic variation using three clusters to represent the seed zones. (c) F_ST_ values between each seed zone at neutral loci (left) and putatively adaptive loci (right).

### 3.4 Sampling to capture wild adaptive variation

Previous estimates of genomic diversity in American chestnut show that the southern portion of the range had the highest levels of heterozygosity, which decreased as latitude increased (Sandercock et al., 2022). As such, we expected that sampling intensity required to capture most adaptive diversity would be lower in the more southern Zone 3, and would increase with each successive latitudinal zone. Consistent with this, we found that fewer trees will need to be sampled from Zone 3, and more trees will need to be obtained from Zone 1 for the 90% and 95% diversity matching targets (Table 2). While the fewest trees will need to be sampled from Zone 3 for the 99% diversity matching target, most trees will instead need to be sampled from Zone 2. Estimates for the central Zone 2 were closer to the Zone 1 estimates across all diversity matching targets (Table 2). An approximately 10-fold difference in sampling intensity will be required to match allele frequencies at 99% compared with 90% coefficients of determination (Table 2). Using a three seed zone model will require more trees to be sampled overall (90%=38.15 trees, 95% = 76.75 trees, 99% = 370.35 trees) compared with the two seed zone model (90%=27.78 trees, 95% = 56.03 trees, 99% = 274.93 trees) or the one seed zone model (90%=26.86 trees, 95% = 55.89 trees, 99% = DNF) (Table 2,Table S4,Table S5).

**Table 2:** Tree sampling estimates to capture adaptive diversity from each American chestnut seed zone.

| Seed zone | % variance explained | # of trees to sample | 95% CI |
| --- | --- | --- | --- |
| 1 | 90% | 14.50 | (13.35, 15.65) |
| 2 | 90% | 13.42 | (12.27, 14.57) |
| 3 | 90% | 10.23 | (9.41, 11.05) |
| 1 | 95% | 28.30 | (26.33, 30.27) |
| 2 | 95% | 27.74 | (25.63, 29.85) |
| 3 | 95% | 20.71 | (19.32, 22.10) |
| 1 | 99% | 132.33 | (124.61, 140.05) |
| 2 | 99% | 136.34 | (127.60, 145.08) |
| 3 | 99% | 101.68 | (94.76, 108.60) |

## 4 Discussion

The goals of this study were to describe the genomic basis of local adaptation in American chestnuts, define seed zones for germplasm conservation, and develop a quantitative framework for sampling recommendations. To do this, we used the SNP dataset generated by Sandercock et al. (2022) to perform genome-wide scans for signatures of adaptation, and to model the partitioning of adaptive variation across the landscape. These analyses revealed polygenic adaptation to climate that can be subdivided into three seed zones from which a relatively small number of trees can be sampled to capture 95% of adaptive diversity.

### 4.1 Climate shapes genomic variation in American chestnut

Like many widespread temperate tree species, climate has significantly shaped genomic variation in American chestnut. Although well connected by gene flow, this species exhibits isolation-by-distance across a natural range with highly variable abiotic environments, in particular precipitation and temperature gradients. The geography of population structure across this range was recently shown to track transitions to mean below freezing winter temperatures in the north, and to a 25% higher annual precipitation in the south (Sandercock et al., 2022). Among *≈*11.5 million SNPs tested, we identified *≈*18.5k associated with climate between RDA and LFMM. The importance of our tested climate variables suggests precipitation, its seasonality, and its interaction with temperature in the summer are primary drivers of adaptation in chestnut, comprising the top three and 11 of the top 15 climate variables in the gradientForest analysis. This contrasts with, for example, lodgepole pine and red spruce, for which variables related to winter temperature are most important to adaptation (Capblancq et al., 2023; Yu et al., 2022). Many of the most studied forest tree species occupy ranges that begin and end further north than that of American chestnut. This dichotomy may reflect a different cold hardiness strategy for chestnut that is insufficient in areas with extremely cold winter temperatures, as well as adaptations to the high summer evaporative demand in the southern portion of its range.

A limitation of both these approaches is that they assume linear relationships between climate and allele frequencies(Capblancq & Forester, 2021; Lotterhos, 2022). Methods to identify nonlinear relationships between genotypes and environment have been developed, such as gradientForest (Fitzpatrick & Keller, 2015), but are not computationally efficient as screening tools for large genomic datasets. Modeling such nonlinear genotype-environment interactions may be of interest in chestnut and other species, particularly given the nonlinear breaks in temperature and precipitation at population boundaries estimated with genome-wide data, and the correspondence of these breaks to seed zones estimated with our adaptive loci.

### 4.2 Adaptive units for germplasm conservation and reforestation

Chestnut restoration efforts will need to account for local adaptation by matching blight resistant families to climatic niches in which they exhibit appropriate timing of growth and dormancy, water use efficiency, drought tolerance, and more. Characterization of the pattern and extent of such local adaptations in forest trees is typically achieved through reciprocal common gardens. As these are not possible for American chestnut, we sought to use correlations between multivariate genotypes to the climate space in which they are found as a proxy for adaptive traits. In both scenarios, the concept of seed zones has been an essential tool for land managers to identify unique regions of climate adaption and to inform candidate areas for reintroduction of seed, often from orchards of known provenance. For chestnut, this matching is further complicated by the hybrid or transgenic nature of the planting stock, which makes provenance a more nebulous concept even if phenotypic local adaptation was well described. Using both genomic and climate information, we found that the American chestnut range can be most parsimoniously divided into three seed zones, which best represents the American chestnut adaptive boundaries. These zones roughly match background population structure in the species (Sandercock et al., 2022), which suggests our putative adaptive cohort of SNPs may be substantially impacted by false positives that simply recapitulate populations structure. We view this as unlikely for three reasons. First, we applied very strict outlier cutoffs, which if anything, biased the dataset toward the exclusion of true positives rather than inclusion of false positives. While the number of adaptive loci in our dataset was very large, it is consistent with the polygenic nature of the traits that usually comprise local adaptation in temperate trees species (Savolainen et al., 2013; Savolainen et al., 2007). Second, F_ST_ for the adaptive SNP cohort was 5.5-6.5-fold higher than the genome-wide background across the seed zones, suggesting divergent selection. Finally, while the three seed zones roughly track background population structure, there were important differences. For example, seed zone one extended southward through high elevation areas of the state of West Virginia, which is not reflected in the neutral population structure map, and there was an extension of Seed Zone 2 southward and Seed Zone 3 northward along the border of Tennessee and North Carolina, which reflects a demarcation between the Great Smokey Mountains in the east (Seed Zone 2) and the much warmer Tennessee Valley to the west (Seed Zone 3).

### 4.3 Matching adaptive portfolios to future climates

American chestnut has clearly adapted to regional precipitation and temperature regimes, and while the above seed zones provide a spatial delineation of how genotypes match geography, these estimates reflect historical climate. Under climate change, the coupling of climate and geography in the context of our modeling will break down, and matching genotypes to planting sites will require reconciling current climates that interact with seedling fitness, and future climates that impinge on competitive ability, reproduction, recruitment, and adaptation of subsequent generations. Our results of areas that are adapted to drier conditions will be expected to have greater maladaptation due to future climate is consistent with previous assessments (Noah et al., 2021). Though, these previous assessments did not include genomic data when estimating the impact of climate change on American chestnut distributions. Nevertheless, genomic offset is predicted to be higher for all areas of the range under the more severe RCP8.5 climate predictions – which is consistent with recent evaluations of American chestnut habitat suitability under climate change conditions (Adeyemo & Granger, 2023).

Climate change will shift the habitable range northward (Noah et al., 2021), thus areas farther north should also be considered for reintroduction. American chestnuts are a slow migrating species – traveling approximately 100m per year after the Last Glacial Maximum (Davis, 2021). As such, assisted migration will be necessary for maladapted regions (Aitken et al., 2008). Chestnuts planted northward outside of the parent range will suffer winter injury, but perform better than other assisted migration species from the eastern United States (Clark et al., 2022). Germplasm from the central and northern seed zones (Seed Zones 2 and 1), exhibits less winter injury and should be considered for use in the most northern replanting locations (Schaberg et al., 2022). We recommend additional evaluations of the three seed zones identified here in common garden and controlled environment experiments to determine how germplasm from each of these seed zones will respond to assisted migration and future climate conditions. The additional information will ensure that locally adapted American chestnut trees will be replanted in regions where they are adapted to thrive in.

### 4.4 Sampling estimates for germplasm conservation of wild adaptive genomic diversity

Ultimately, we sought to estimate the number of trees to sample from each seed zone to capture most of the adaptive diversity present in the wild. These sampled trees will be used for germplasm conservation, and to introgress adaptive variation into disease-resistant populations. To make these sampling recommendations, we developed a novel computational method since an existing tool/pipeline was not available. Consistent across the three diversity percentages, 90%, 95%, and 99%, we found that the fewest amount of trees will need to be sampled from the southern Seed Zone 3, and the most from the northern Seed Zone 1.

These findings were expected considering that genomic diversity is highest in the southern chestnut range and decreases as latitude increases (Gailing & Nelson, 2017; Müller et al., 2018; Sandercock et al., 2022; Spriggs & Fertakos, 2021). Thus, fewer trees would need to be sampled from regions of higher heterozygosity and nucleotide diversity to capture the desired percentage of adaptive diversity.

### 4.5 Conclusions

The American chestnut is a functionally extinct species facing both biotic and abiotic threats. Developing disease-resistant chestnut populations is an important first step, but these populations require sufficient adaptive variation to survive and thrive in their replanting locations. However, information regarding the adaptive genomic landscape of the American chestnut was lacking and methods were not available to estimate the number of trees necessary to sample from wild populations to inform germplasm and breeding plans. Therefore, the seed zones identified here, which represent areas of homogeneity relative to the chestnut adaptive genotypes, will be used as sources of germplasm and regions for reintroduction. We found that 18,483 SNPs were significantly associated with climate, that three seed zones best separates the chestnut range into unique regions of adaptive diversity, and that fewer trees will need to be sampled from the southern seed zones to capture the same proportion of adaptive diversity as the more northern seed zones – likely due to the more heterozygosity present in the southern portion of the chestnut range. Future work will need to estimate genomic diversity already present in the backcross breeding program, and to develop a breeding strategy to introgress additional adaptive diversity from the subsampled individuals into the backcross program. Additionally, evaluations should be done to estimate candidate regions for reintroduction of these locally adapted, disease-resistant populations to survive current and future climate conditions. Future breeding simulations will be performed which utilize the sampling estimates and adaptive genomic dataset developed here – leading to the development of locally adapted and disease-resistant populations for the restoration of the American chestnut.

## Acknowledgements

We would like to thank TACF volunteers who participated in sampling of wild trees for this project; Jamie Van Clief (TACF), Loren Hostetter (ACCF), Warren Nance (U.S. Forest Service), and Chance Parker (U.S. Forest Service) for assistance in additional leaf collections; and Advanced Research Computing at Virginia Tech for providing computational resources. This research was supported by the United States National Institute for Food and Agriculture Projects 1018599, 1027966, 1025004 and by a Graduate Fellowship to AS from the Virginia Tech Institute for Critical Technology and Applied Science.

## Contributions

A.M.S, J.W.W, and J.A.H designed the study; A.M.S. and J.A.H. analyzed the data; QZ contributed to data collection; A.M.S., J.W.W., and J.A.H. wrote the manuscript.

## Data Accessibility

The script for the Python sampling method can be found at https://github.com/alex-sandercock/Capturing_genomic_diversity

