## Supplemental Tables and Figures for "The road to restoration: Identifying and conserving the adaptive legacy of American chestnut"

Table S1: The 32 climate variables from ClimateNA that were used for this study and their respective descriptions.

| Abbreviation | Description |
|---|---|
| Annual variables directly calculated from monthly variables | |
| MAT | mean annual temperature (°C) |
| MWMT | mean warmest month temperature (°C) |
| MCMT | mean coldest month temperature (°C) |
| TD | temperature difference between MWMT and MCMT, or continentality (°C) |
| MAP | mean annual precipitation (mm) |
| AHM | annual heat-moisture index ((MAT+10)/(MAP/1000)) |
| SHM | summer heat-moisture index ((MWMT)/(MSP/1000)) |
| Annual variables derived from monthly variables | |
| DD_0 | degree-days below 0°C, chilling degree-days |
| DD5 | degree-days above 5°C, growing degree-days |
| DD_18 | degree-days below 18°C, cooling degree-days |
| DD18 | degree-days above 18°C, heating degree-days |
| NFFD | the number of frost-free days |
| FFP | frost-free period |
| bFFP | the day of the year on which FFP begins |
| eFFP | the day of the year on which FFP ends |
| PAS | precipitation as snow (mm). For individual years, it covers the period between August in the previous year and July in the current year |
| EMT | extreme minimum temperature over 30 years (°C) |
| EXT | extreme maximum temperature over 30 years (°C) |
| Eref | Hargreaves reference evaporation (mm) |
| CMD | Hargreaves climatic moisture deficit (mm) |
| MAR | mean annual solar radiation (MJ m-2 d-1) |
| RH | mean annual relative humidity (%) |
| CMI | Hogg's climate moisture index (mm) |
| DD1040 | degree-days above 10°C and below 40°C |
| Seasonal variables directly calculated from monthly variables | |
| Tave_wt | winter mean temperature (°C) |
| Tave_sp | spring mean temperature (°C) |
| Tave_sm | summer mean temperature (°C) |
| Tave_at | autumn mean temperature (°C) |
| PPT_wt | winter precipitation (mm) |
| PPT_sp | spring precipitation (mm) |
| PPT_sm | summer precipitation (mm) |
| PPT_at | autumn precipitation (mm) |

Table S2: The environmental variable loadings on the three PC axes used for the LFMM analysis.

| | PC1 | PC2 | PC3 |
|---|---|---|---|
| RH | -0.048 | -0.034 | 0.937 |
| Eref | 0.378 | 0.324 | -0.111 |
| PPT_wt | 0.416 | -0.116 | 0.053 |
| EXT | 0.039 | 0.472 | -0.073 |
| TD | -0.437 | 0.01 | -0.134 |
| MAR | 0.443 | 0.006 | 0.027 |
| PPT_sm | 0.351 | -0.256 | -0.156 |
| CMI | 0.132 | -0.498 | 0.111 |
| AHM | -0.119 | 0.487 | 0.156 |
| DD_18 | -0.373 | -0.326 | -0.148 |

Table S3: Number of single-nucleotide polymorphisms (SNPs) in coding and noncoding features. Three datasets were evaluated: **Adaptive** (18,483 putatively adaptive SNPs from the GEA analyses), **Random** (randomly selected SNPs with allele frequency distribution matched to the adaptive set), and **Full** (original filtered SNP dataset from Sandercock et al. (2022) with MAF<0.05 filter). The promoter region includes the 2kb region upstream of the mRNA intervals.

| | Adaptive | Random | Full |
|---|---|---|---|
| Number of SNPs in CDS | 264 | 619 | 377161 |
| Number of SNPs in exon | 435 | 991 | 611402 |
| Number of SNPs in five_prime_UTR | 69 | 141 | 89577 |
| Number of SNPs in three_prime_UTR | 104 | 236 | 150477 |
| Number of SNPs in mRNA | 2120 | 3462 | 2102039 |
| Number of SNPs in promoter | 1309 | 1830 | 1154261 |
| Number of SNPs in gene | 2120 | 3462 | 2102039 |
| Number of SNPs in intron | 1712 | 2537 | 1528332 |
| Total SNPs in dataset | 18483 | 18483 | 11526713 |

Table S4: Estimated number of samples to capture adaptive diversity from each seed zone for a two seed zone model.

| Seed zone | % variance explained | # of trees to sample | 95% CI |
|---|---|---|---|
| 1 | 90% | 16.73 | (15.69, 17.77) |
| 2 | 90% | 11.05 | (10.13, 11.97) |
| 1 | 95% | 34.12 | (31.90, 36.34) |
| 2 | 95% | 21.91 | (20.11, 23.71) |
| 1 | 99% | 167.56 | (156.37, 178.75) |
| 2 | 99% | 107.37 | (100.20, 114.54) |

Table S5: Estimated number of samples to capture adaptive diversity under a single seed zone model. The estimate for the 99% variance explained did not complete due to excessive run time.

| Seed zone | % variance explained | # of trees to sample | 95% CI |
|---|---|---|---|
| 1 | 90% | 26.86 | (24.55, 29.17) |
| 1 | 95% | 55.89 | (51.01, 60.77) |
| 1 | 99% | DNF | DNF |

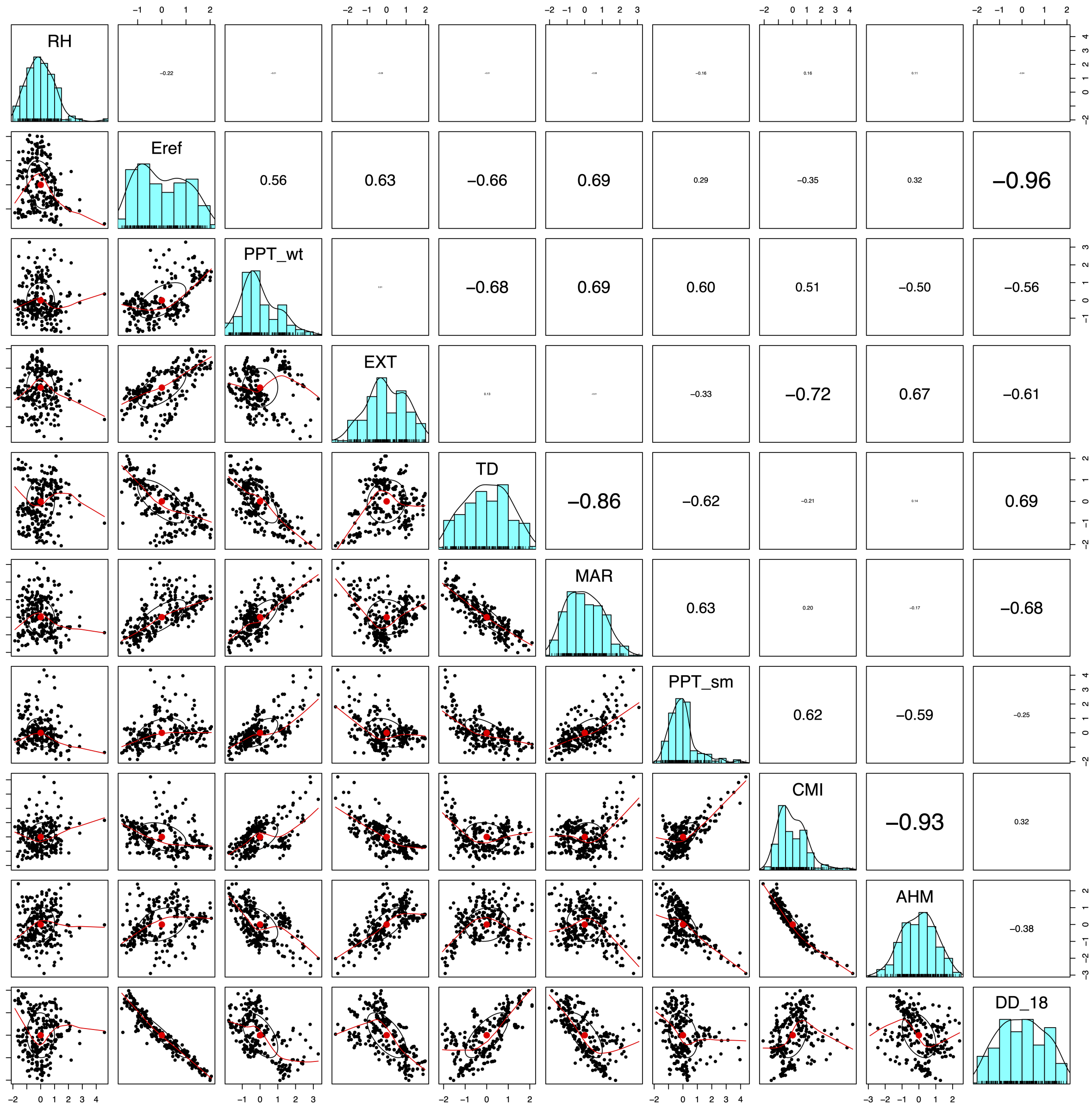


Figure S1: Ten uncorrelated climate variables were selected for use in the genotype-environment association analyses. Pairs plot of the selected 10 environmental variables. The bottom half are regression plots and the top half are Pearson's correlations.

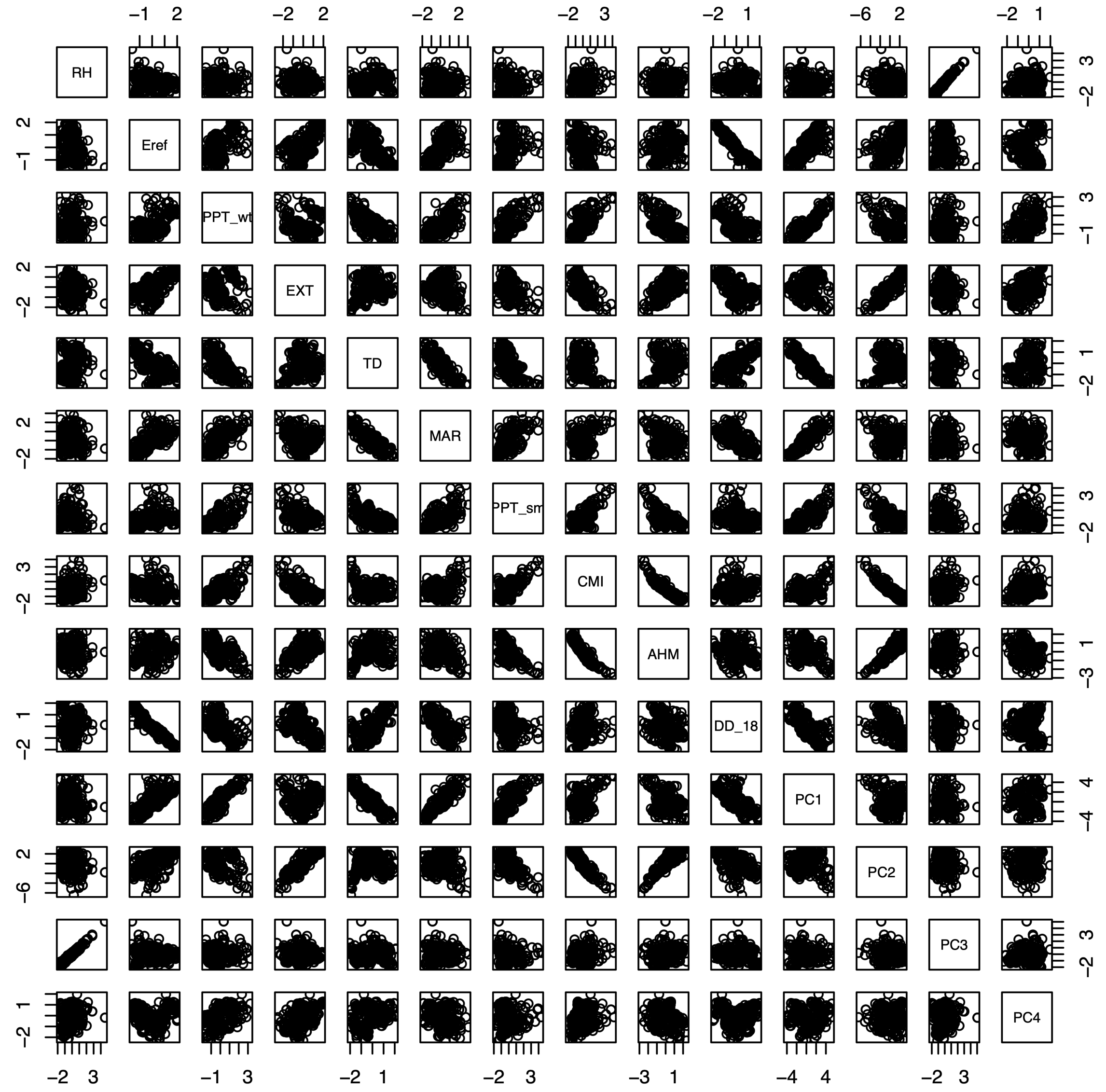


Figure S2: Three PC axes were used as synthetic variables for the LFMM genotype-environment association analyses. Pairs plot of the selected 10 environmental variables and the first four PCs from the PCA of the 10 climate variables. The bottom half and top half are linear regressions between the variables.

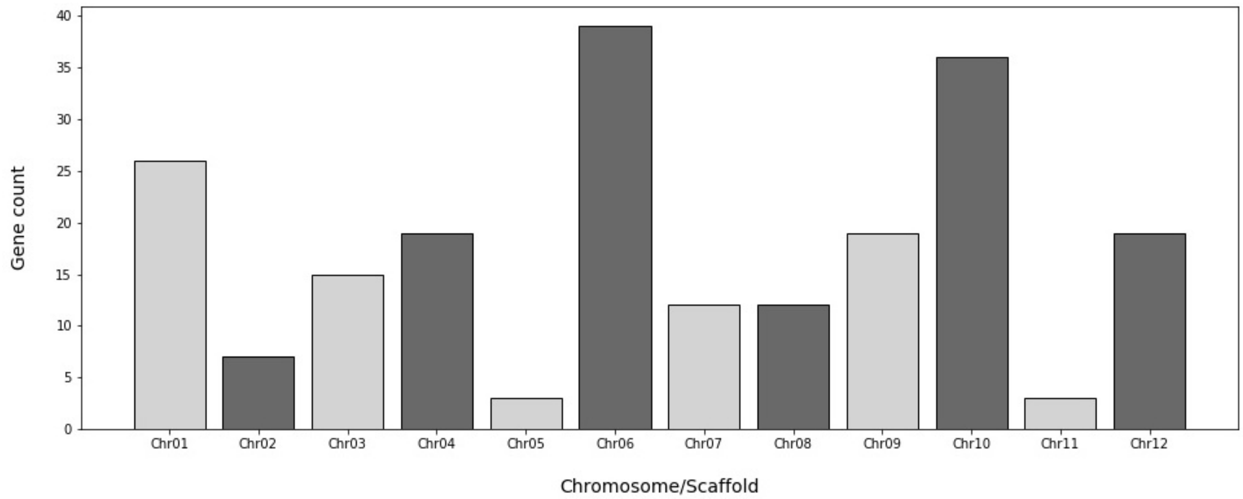


Figure S3: The number of genes that contained at least one adaptive loci within each *Castanea dentata* chromosome. Scaffolds that did not contain genes with adaptive loci were omitted from the figure.

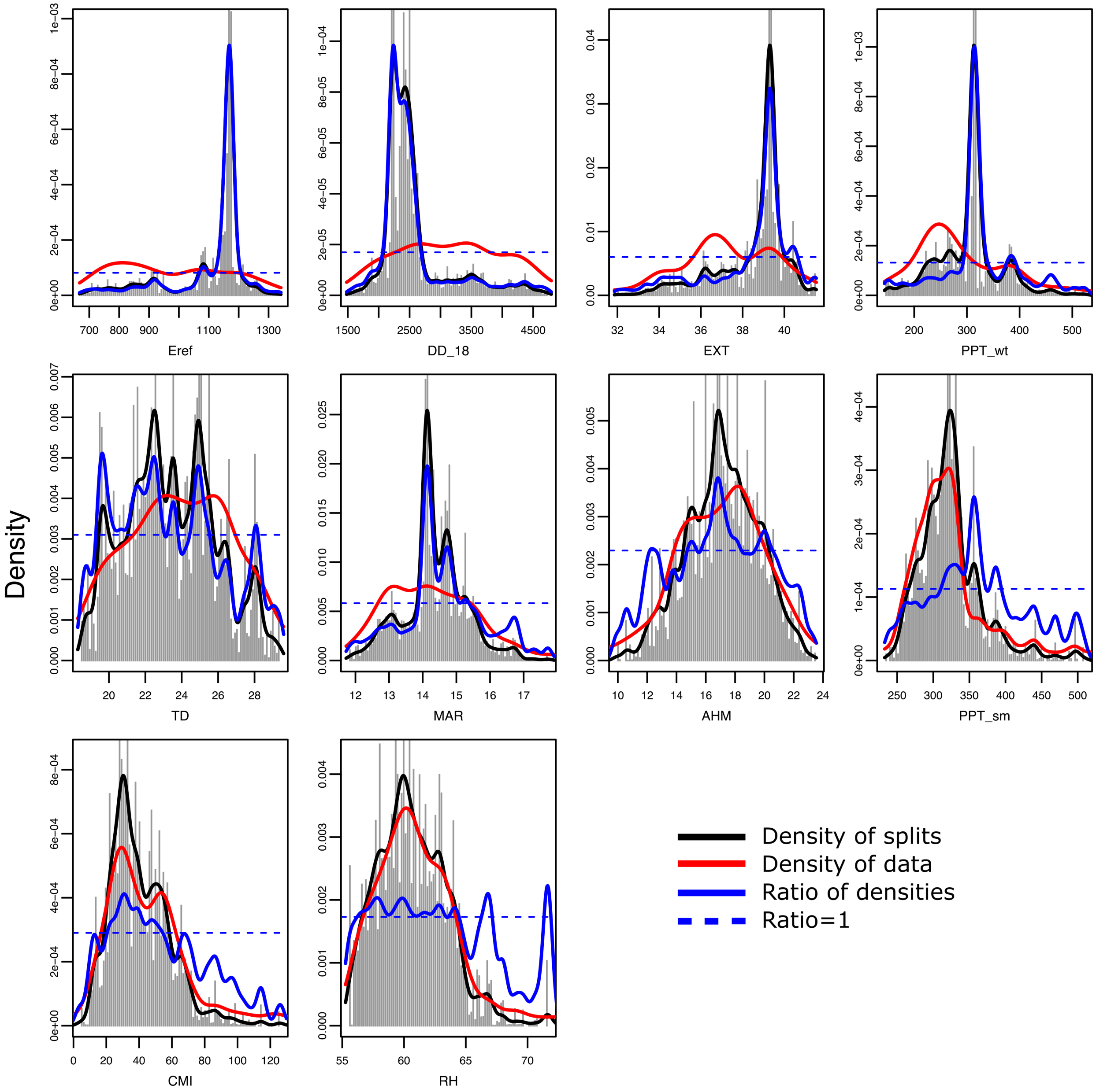


Figure S4: Gradient Forest adaptive allelic turnover for each of the ten climate variables and 18,483 putatively adaptive loci. Spikes in the binned split importance reveal where changes in adaptive allele density are occurring along each climate gradient.

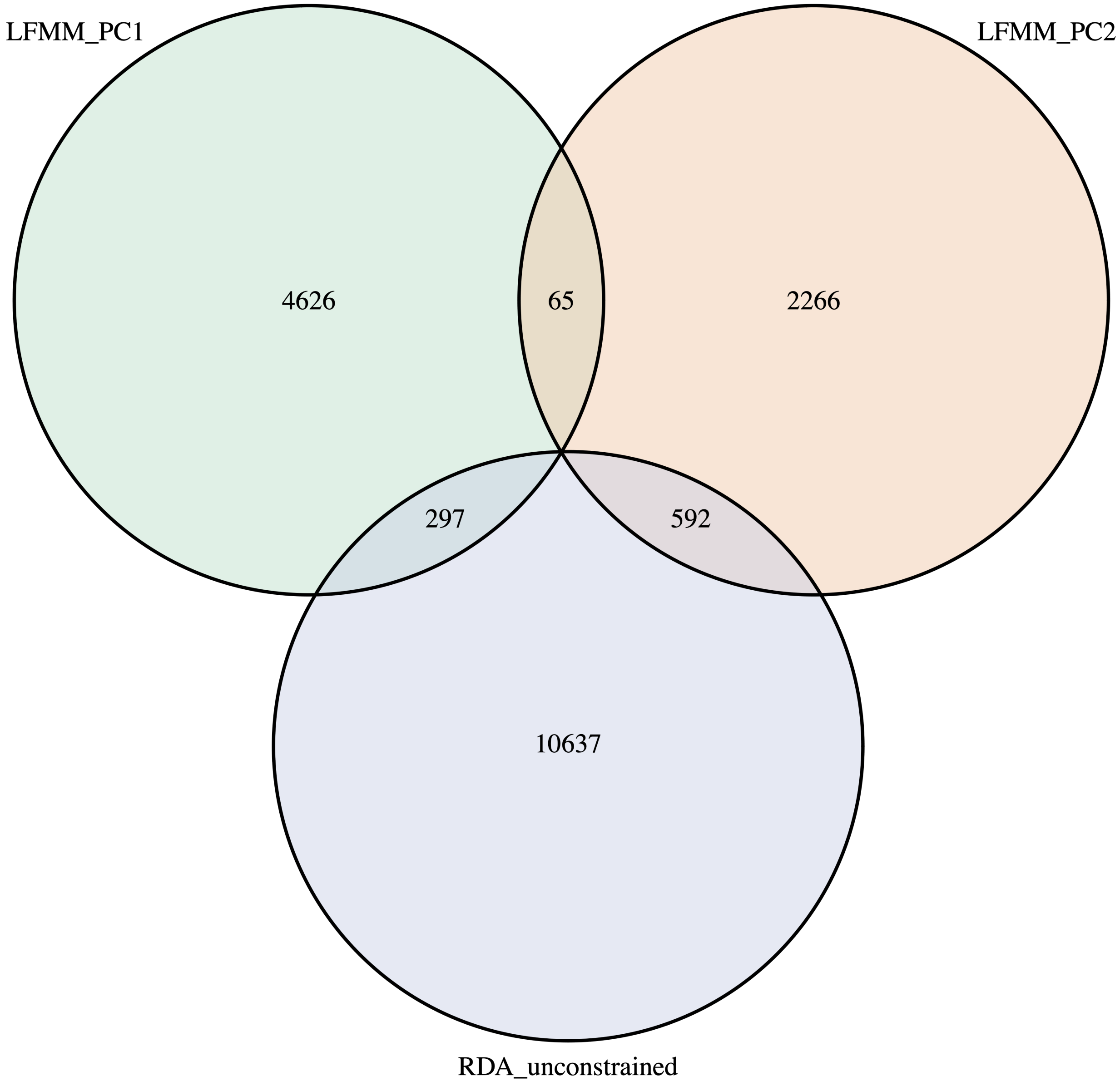


Figure S5: Venn diagram for the 18,483 adaptive loci identified using RDA and LFMM. For LFMM, only the SNPs from the PC1 and PC2 analyses are shown due to the PC3 analysis finding zero outlier loci.

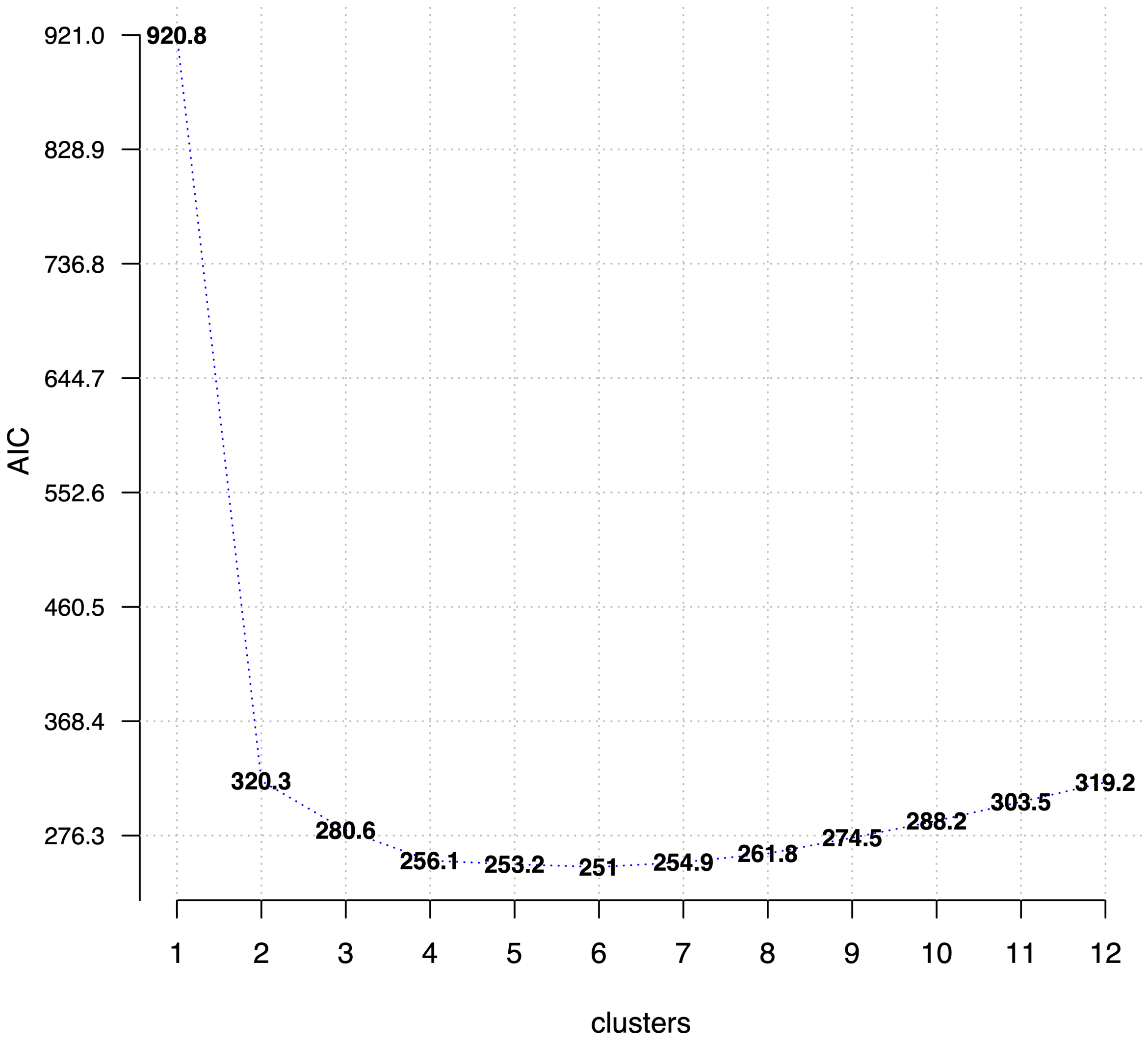


Figure S6: AIC score evaluation to determine optimal number of seed zones.

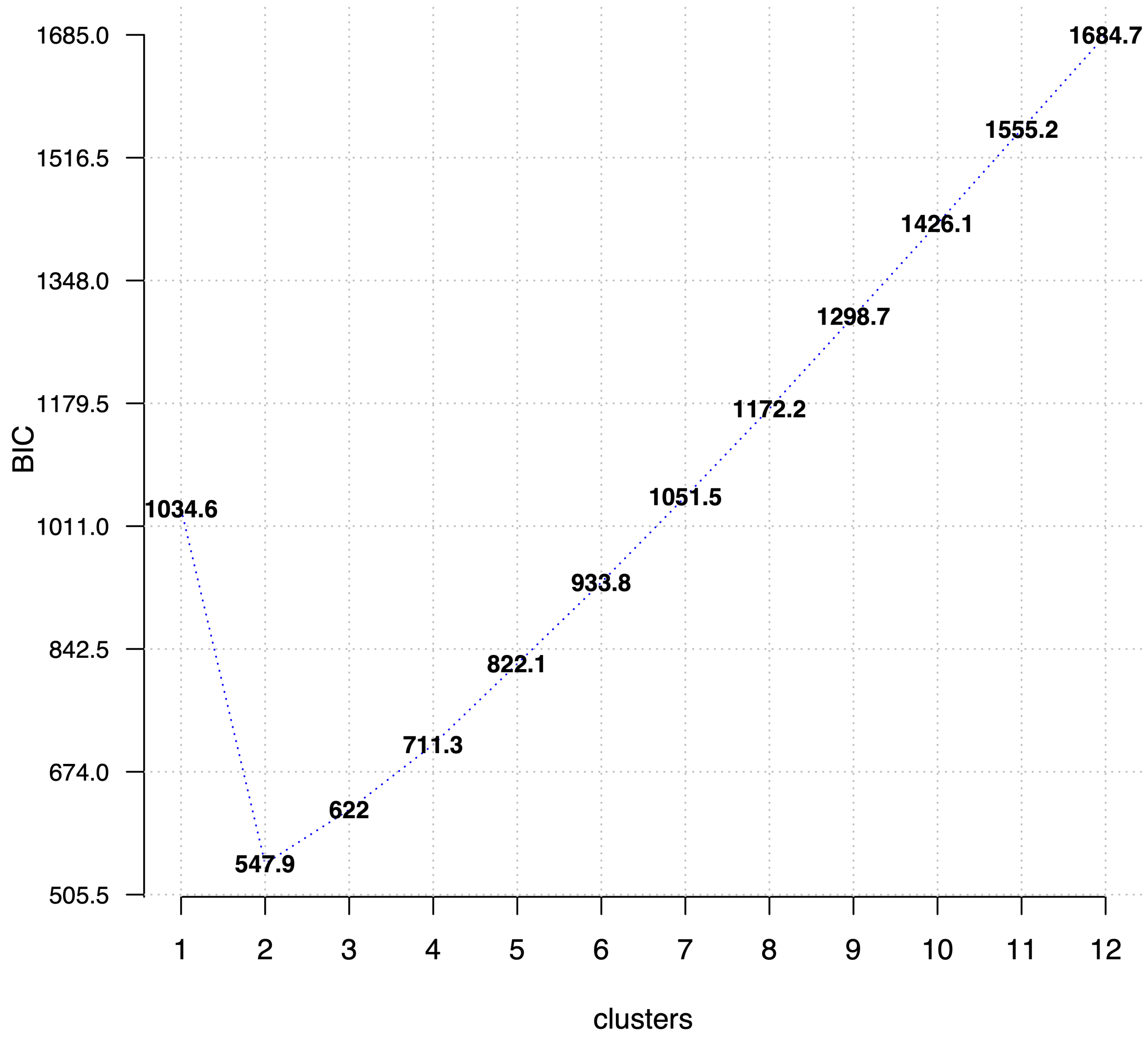


Figure S7: BIC score evaluation to determine optimal number of seed zones.

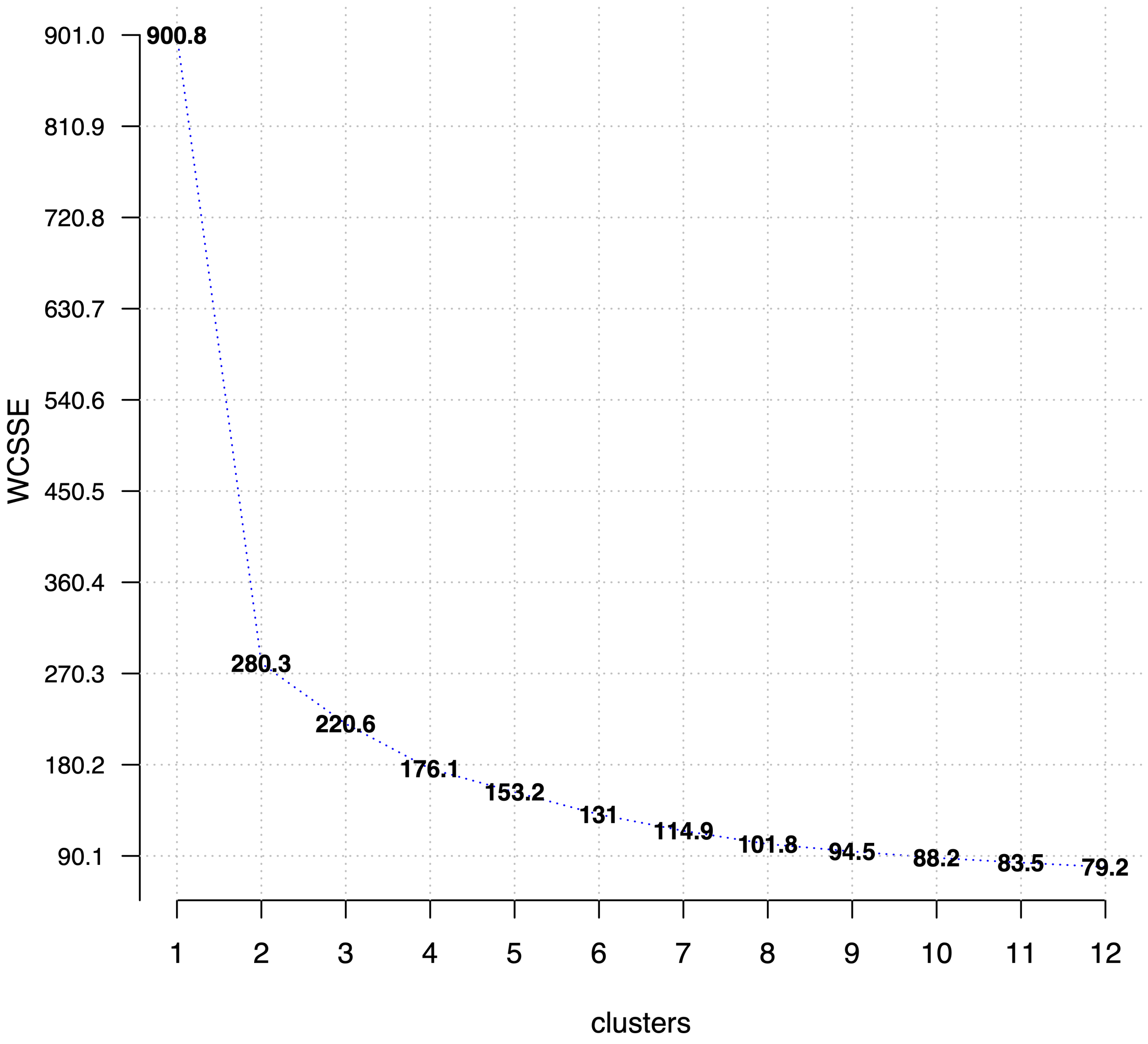


Figure S8: WCSSE score evaluation to determine optimal number of seed zones.

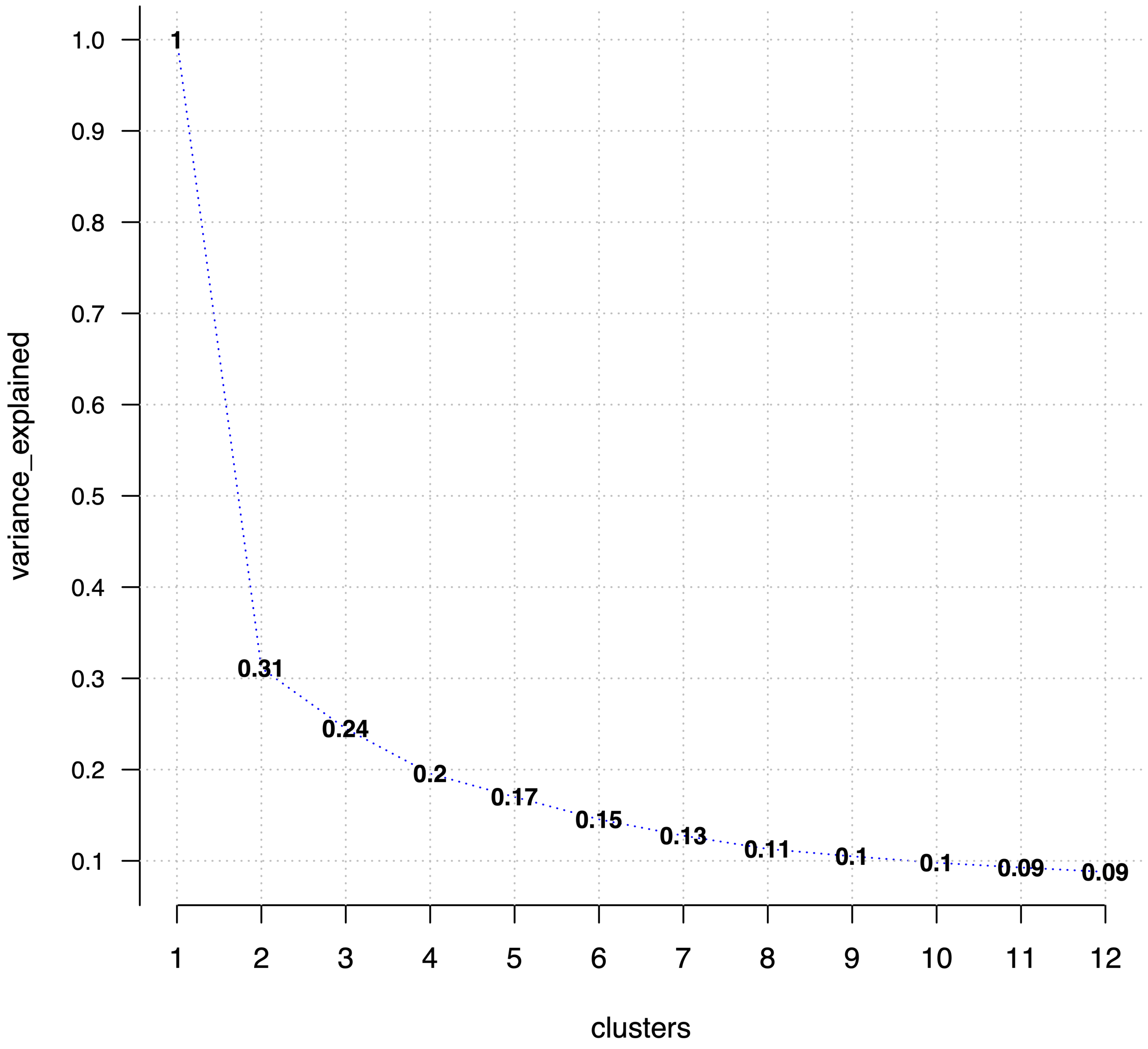


Figure S9: Variance explained scores to determine optimal number of seed zones.
